# Exportin 1 is required for the reproduction and maize mosaic virus accumulation in its insect vector *Peregrinus maidis*

**DOI:** 10.1101/2023.09.19.558515

**Authors:** Cesar A. D. Xavier, Clara Tyson, Leo M. Kerner, Anna E. Whitfield

## Abstract

Exportin 1 (XPO1) is the major karyopherin-β nuclear receptor mediating the nuclear export of hundreds of proteins and some classes of RNA and regulates several critical processes in the cell, including but not limited to, cell-cycle progression, transcription, translation, oncogenesis and longevity. Viruses have co-opted XPO1 to promote nucleocytoplasmic transport of viral proteins and RNA. Maize mosaic virus (MMV) is an *Alphanucleorhabdovirus* transmitted in a circulative propagative manner by the corn planthopper, *Peregrinus maidis*. MMV replicates in the nucleus of plant and insect hosts, and it remains unknown whether MMV co-opts *P. maidis XPO1* (*PmXPO1*) to complete its life cycle. Because XPO1 plays multiple regulatory roles in cell functions and virus infection, we hypothesized that RNAi-mediated silencing of *XPO1* would simultaneously and negatively affect MMV accumulation and insect physiology. Although *PmXPO1* expression was not modulated during MMV infection, *PmXPO1* knockdown negatively affected MMV accumulation in *P. maidis* at 12 and 15 days after microinjection. Likewise, *PmXPO1* knockdown negatively affected *P. maidis* survival and reproduction. *PmXPO1* exhibited tissue specific expression patterns with higher expression in the ovaries compared to the guts of adult females. Survival rate was significantly lower for *PmXPO1* knockdown females, compared to controls, but no effect was observed for males. Adult females with *PmXPO1* knockdown were heavier and had a larger abdomen compared to controls at 4, 8 and 12 days after dsRNA microinjection. Consistent with an increase in weight, glyceride content specifically and significantly increased in *PmXPO1* knockdown female planthoppers. Ovary development was significantly inhibited, and mature eggs were not observed in adult females with *PmXPO1* knockdown. Consistent with a major role of *Pm*XPO1 in ovary function and egg production, oviposition and egg hatch in plants was dramatically reduced in dsRNA *PmXPO1* treated insects compared with control. Altogether, these results suggest that *PmXPO1* is a positive regulator of *P. maidis* reproduction and that it plays a proviral role in the insect vector supporting MMV infection.

## INTRODUCTION

In eukaryotic cells, the evolution of nuclear compartmentalization of the genome from other cellular components was followed by the emergence of a tightly regulated nucleocytoplasmic transport machinery of proteins and RNAs that plays a vital regulatory role in physiology and cell development (Wing et al., 2022, Terry et al., 2007). Nuclear pore complexes (NPC) are the main gates through which molecules and macromolecular complexes traffic from or into the nucleus providing a selective and regulated movement of macromolecules (Knockenhauer and Schwartz, 2016). While small soluble molecules can passively diffuse through NPC, the bidirectional movement of molecules larger than 40 kDa is actively regulated and requires the specific action of nuclear transport receptors. The Karyopherin-β (Kap-β) family consists of the main group of nuclear receptors mediating nucleocytoplasmic transport and several classes of Kap-β receptors involved in import (importins), export (exportins) and bidirectional transport (biporins) of cargoes have been characterized and demonstrated to be conserved across the main groups of eukaryotes (Wing et al., 2022, O’Reilly et al., 2011). Exportin 1 (XPO1, also known as CRM1) is the major Kap-β nuclear receptor mediating the nuclear export of hundreds of proteins and some classes of RNAs via adapter proteins, including viral/cellular mRNA, ribosomal RNA, and small nuclear RNAs (Watanabe et al., 1999, Stade et al., 1997, Fornerod et al., 1997, Fischer et al., 1995). By selectively binding to leucine-rich nuclear export signal (NES) on the cargo protein, along with Ran-GTP, XPO1 exports and controls nuclear/cytoplasmic distribution of several cellular factors and regulates crucial processes in the cell, including cell-cycle progression, transcription, translation, oncogenesis, longevity, and immune response (Kırlı et al., 2015, Fornerod et al., 1997). Interfering with XPO1 activity can significantly affect cell physiology and the potential use of small molecules selectively and reversibly inhibiting XPO1 has been explored as a therapeutic approach to treat some types of cancer and more recently virus infection (Azmi et al., 2021, Uddin et al., 2020, Azizian and Li, 2020).

Numerous viruses have co-opted nucleocytoplasmic machinery to promote transport of viral proteins and RNAs. Since the majority of viral proteins are multifunctional, playing distinct roles in the nucleus-cytoplasm interface, tight regulation of protein distribution is critical for virus infection and several functional nuclear localization signals (NLS) and NES signals have been characterized in viral-encoded proteins (Zhang et al., 2022, Yang et al., 2022, Cao et al., 2022, Huang et al., 2013, Ghildyal et al., 2009, Fischer et al., 1995). XPO1 is the main host factor mediating nuclear export of viral proteins and ribonucleoprotein (vRNP) complex and its role in nuclear and cytoplasmic virus infection has been demonstrated (Yang et al., 2022, Zhang et al., 2021, Anderson et al., 2018, Boons et al., 2015, Huang et al., 2013, Neville et al., 1997). Direct interaction of viral proteins and XPO1 has been characterized for a diverse group of RNA and DNA viruses infecting vertebrates, including influenza virus, Hepatitis B virus (HBV), respiratory syncytial virus (RSV) and human immunodeficiency virus (HIV; Yang et al., 2022, Su et al., 2022, Huang et al., 2013, Ghildyal et al., 2009, Elton et al., 2001, Neville et al., 1997, Fischer et al., 1995). In all cases, selective inhibition of XPO1 activity significantly impaired virus life cycle and has been explored as a potential prophylactic approach to control virus infection in mammalian cells (Mathew et al., 2021, Uddin et al., 2020, Mathew and Ghildyal, 2017). Although far less studied in plant viruses, NES-containing viral proteins, and their interaction with XPO1 have been characterized. For example, XPO1 mediates the nuclear export of sumoylated RNA-dependent RNA polymerase (RdRp) of turnip mosaic virus (TuMV), a cytoplasmic RNA virus of the genera *Potyvirus*, and XPO1 silencing negatively affected TuMV replication and accumulation in *Arabidopsis thaliana* and *Nicotiana benthamiana* hosts (Zhang et al., 2022, Zhang et al., 2021). Moreover, a NES was identified in the matrix protein of potato yellow dwarf virus (PYDV), a plant rhabdovirus that replicates in the nucleus of infected cells, and its interaction with *A. thaliana* XPO1 was demonstrated (Anderson et al., 2018). However, the significance of this interaction within the virus life cycle remains unclear and the mechanisms involved in regulation of nucleocytoplasmic trafficking of proteins and RNAs for nuclear-replicating plant rhabdoviruses remains poorly understood.

Despite the well-established proviral role of XPO1 during virus infection in vertebrate and plant hosts, a general antiviral response has been reported against arthropod-borne viruses (arboviruses) including flavivirus, togavirus, bunyavirus and rhabdovirus (Yasunaga et al., 2014). Knockdown of XPO1 in whole body *Drosophila* led to increased replication of West Nile virus (WNV) and vesicular stomatitis virus (VSV) which in turn negatively impacted insect survival (Yasunaga et al., 2014). An XPO1 antiviral response was also demonstrated in mammalian cells, suggesting this protein has a conserved role across distantly related hosts that support arbovirus infection (Yasunaga et al., 2014). Although the viruses in these studies replicate in the cytoplasm, nucleocytoplasmic movement of proteins is essential for virus replication and interaction between viral protein and XPO1 has been characterized for flaviviruses (De Jesús-González et al., 2023, Pryor et al., 2006, Rawlinson et al., 2009) and a rhabdovirus (Pasdeloup et al., 2005). How viruses manage to overcome the trade-off between pro- and antiviral XPO1 activity has to be further investigated. While exploring XPO1 as a target to control arbovirus transmission is potentially promising, either by directly impairing virus replication or indirectly affecting survival and physiology of the insect vector, characterizing the impact of XPO1 knockdown on the physiology of arboviruses vectors is still an essential prerequisite. Most of the studies investigating the role of XPO1 in insects physiology were based on the *Drosophila* model. Because XPO1 plays pleiotropic roles and its inhibition negatively affects cell function, studies investigating the physiological roles of XPO1 during virus infection have mostly been performed in cell cultures and the effect of its knockdown in whole body of arbovirus vectors is not well known.

Maize mosaic virus (MMV) is an *Alphanucleorhabdovirus* (family *Rhabdoviridae*) transmitted in a circulative propagative manner by the corn planthopper, *Peregrinus maidis* (Dietzgen et al., 2020, Hogenhout et al., 2008). MMV replicates in the nucleus of plant and insect hosts and persists throughout the entire insect life cycle (Barandoc-Alviar et al., 2016, Ammar el et al., 2009). Localization of MMV proteins is conserved in plant and insect cells, suggesting a similar replication strategy in both hosts (Martin and Whitfield 2018). However, it is still unknown how viral proteins, transcripts and vRNPs traffic from or into the nucleus during virus replication in the insect vector. We hypothesized that MMV co-opts *P. maidis XPO1* (*PmXPO1*)-encoded protein to promote nucleocytoplasmic traffic of viral proteins or vRNPs and silencing of *PmXPO1* would negatively affect virus accumulation. To address this hypothesis, we studied the effect of *PmXPO1* knockdown on MMV accumulation in whole bodies of male and female insects. Moreover, we were interested in understanding the effect of *PmXPO1* knockdown in the physiology of the planthopper vector. Because XPO1 plays multiple regulatory roles in cell functions, we hypothesized that XPO1 knockdown would significantly impair *P. maidis* physiology. By using an *in vivo* RNA interference (RNAi) approach, this study provides the first evidence that *PmXPO1*-encoded protein plays a proviral role in the insect vector supporting MMV infection and that it is a positive regulator of *P. maidis* reproduction.

## RESULTS AND DISCUSSION

### Identification and molecular characterization of *PmXPO1*

To identify *PmXPO1* coding sequence, a local BLASTn search was performed in two previously independently assembled *P. maidis* transcriptomes, which analyzed the transcriptional landscape across all developmental stages (Wang et al., 2023b) and the response to virus infection in whole-body of adult individuals (Martin and Whitfield, 2018). Using the *Drosophila melanogaster XPO1* (*DmXPO1*) nucleotide sequence as a query, we identified one transcript from each data set homologous to the *DmXPO1* coding sequence (**Supp. File 1**). The transcripts shared 99.61% of nucleotide identity between them and were predicted to encode an open reading frame (ORF) of 3,216 base pairs (bp), which is in the range of ORFs encoding XPO1 protein from other organisms (**Supp. Table S2**). Analysis of cloned sequences further confirmed the authenticity of the *in silico* identified transcripts. They were predicted to encode an ORF with exactly 3,216 bp that shared 99.71 to 100% of amino acid identity with the *in silico* identified transcripts. A domain-based blast search performed against the Conserved Domain Database (CDD) identified the typical CRM1 importin beta-related nuclear transport receptor domain superfamily (accession COG5101). Furthermore, domain architecture analysis revealed the presence of three conserved domain characteristics of XPO1 proteins (IBN-N, XPO1, CRM1_C; E-value < 3.95e-9; **Figure 1**). Further sequence comparisons demonstrated that PmXPO1 is closely related to the predicted XPO1 sequence from the brown planthopper, *Nilaparvata lugens* (locus LOC111047279; 97.57% of aa identity), and highly conserved at amino acid level with orthologs from a diverse group of insects including those from orders Hemiptera, Blattodea, Diptera, Hymenoptera and Orthoptera (**Figure 1**).

**Figure 1.**
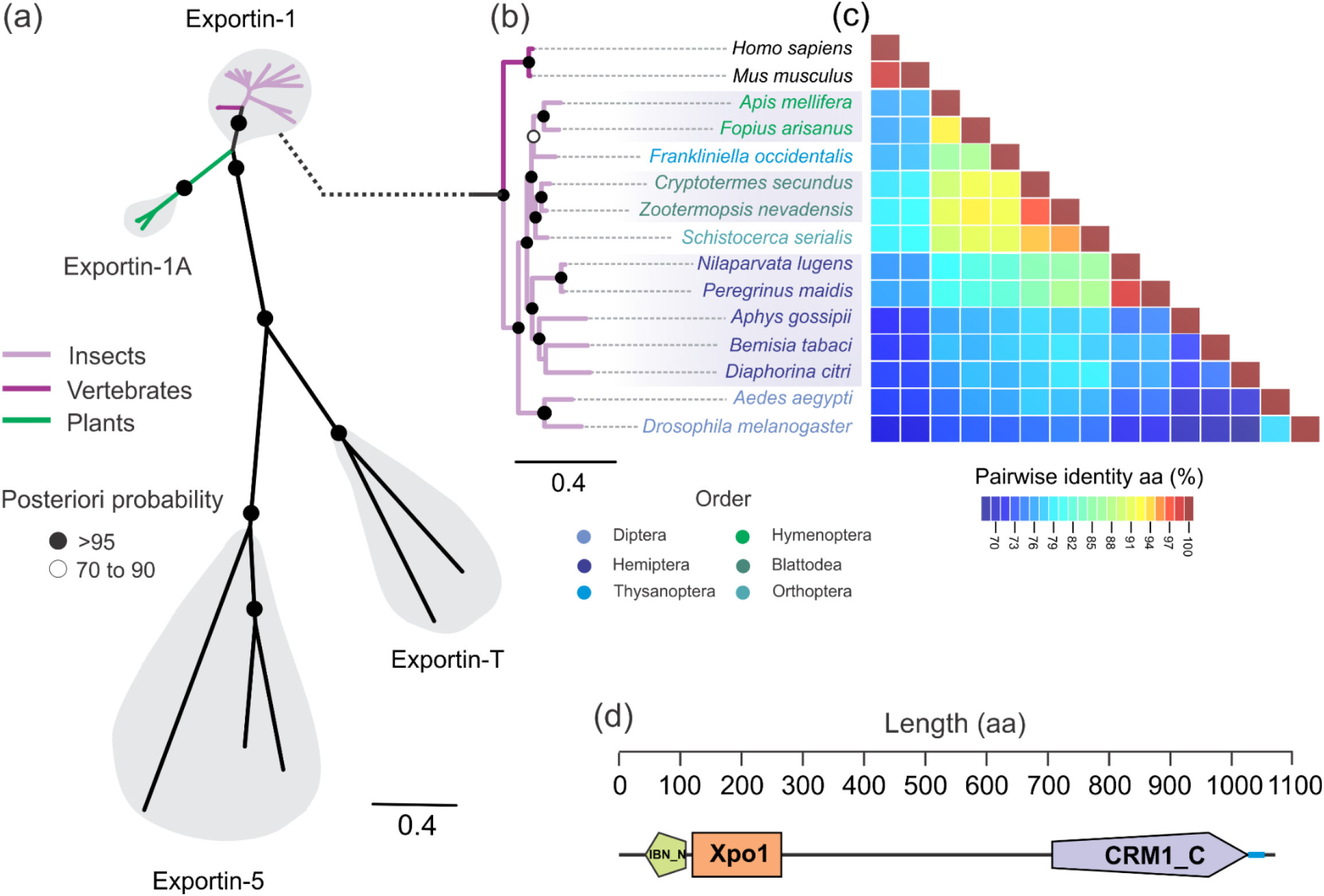
Identification and molecular characterization of *P. maidis exportin 1* (*PmXPO1*). (**a**) Unrooted Bayesian phylogenetic tree constructed with the full-length amino-acid sequences of the PmXPO1 and representative sequences from species of the orders Blattodea, Diptera, Hemiptera, Hymenoptera, Orthoptera, and Thysanoptera. Representative sequences of other exportins (exportin-1A, exportin-5 and exportin-T) from karyopherin-β subfamily closest related to XPO1 were added into the phylogeny (for further sequences information see **Supp. Table S2**). **(b)** Magnification of the clade that includes XPO1 sequences from arthropods and vertebrates. Species names of insects are colored according to the order, as indicated under the phylogeny. **(c)** Pairwise identity matrix generated using XPO1 amino acid sequences. Colors indicate the identity levels shown in the scale bar under the matrix. (**d**) Domain architecture of PmXPO1 showing the three canonical XPO1 domains: Importin-beta N-terminal domain (IBN-N, accession number: SM000913, *E* = 3.95e-9), exportin 1 (XPO1, accession number: PF08389, *E* = 7.4e-42) and CRM1 C terminal (CRM1_C, accession number: SM001102; *E* = 3.24e-183).

To further validate the identified sequence and to investigate the evolutionary relationship between *PmXPO1* and those from other insects, a phylogenetic tree based on the full-length XPO1 amino acid sequence was constructed using Bayesian inference. Overall, XPO1 sequences representative of six insects orders and two sequences from vertebrates (*Homo sapiens* and *Mus musculus*) were well conserved at amino acid level, as indicated by the short length of internal nodes in the phylogenetic tree and identity analysis (**Figure 1**). Insect XPO1 sequences clustered in a monophyletic clade which segregated according to insect order (**Figure 1**). Specifically, the PmXPO1 clustered in a clade composed of hemipterans in a branch closest related to *N. lugens* (**Figure 1**). The use of different sequence analysis approaches enabled the reliable identification and characterization of *PmXPO1*, the first step towards the functional characterization of this gene in *P. maidis*.

### Expression analyses of *PmXPO1*

To understand the biological functions that XPO1 may play in the corn planthopper, we first evaluated gene expression across developmental stages and female tissues, ovary and gut, using reverse transcription quantitative real time PCR (RT-qPCR). *PmXPO1* were expressed at all developmental stages and tissues analyzed (**Figure 2a, b**). Analysis of *PmXPO1* expression in whole bodies of insects across life stages, nymphs (N1 to N5) and adults (males and females), revealed no significant difference in transcript abundance (**Figure 2a**). In contrast, *PmXPO1* exhibited tissue specific expression patterns with a 2.6-fold higher expression in the ovaries compared to the guts of adult females (**Figure 2b**). The very conserved role of XPO1 on mediating nuclear transport of thousands of proteins and some classes of RNAs, which is essential for maintaining the cellular homeostasis, most likely makes this protein equally important across all developmental stages not showing any specific expression patterns at transcriptional level. However, the significantly higher transcript abundance observed in ovaries indicates that XPO1 may play a role in regulating ovary function and consequently reproduction.

**Figure 2.**
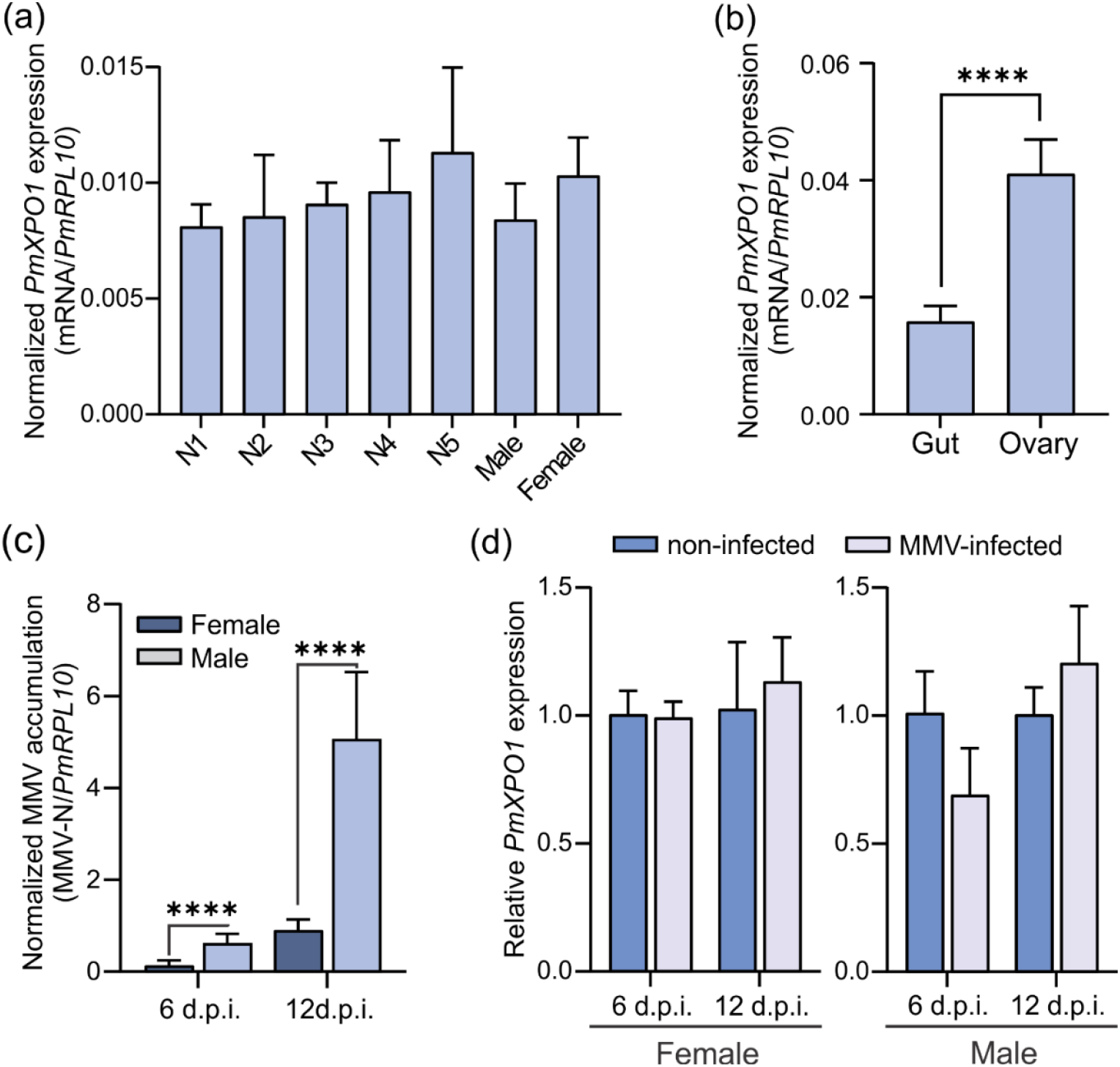
Expression analysis of *P. maidis exportin 1* (*PmXPO1*) according to developmental stages and MMV infection status. (**a**) *PmXPO1* expression was analysed across developmental stages including nymphs (N1, N2, N3, N4 and N5) and adults (males and females). Bars indicate mean and error bars standard deviation of five biological replicates composed by a variable number of individuals according to the stage (N1 = 20, N2 and N3 = 10, N4 and N5 = 5, male and female = 3). No significant difference among treatments was found by one-way ANOVA. (**b**) *PmXPO1* expression differs between the gut and ovary tissues of female planthoppers. Bars indicate mean and error bars standard deviation of five biological replicates composed by five and six dissected ovaries and guts, respectively. Significant differences between groups were assessed by t-test (*** *p*<0.001). (**c**) Viral titer measured at 6- and 12-days post-injection (d.p.i.) and compared between males and females on the samples used in **d.** (**d**) *PmXPO1* expression was not modulated during maize mosaic virus (MMV) infection at 6 and 12 days after virus injection in males and females. In **c** and **d** bars indicate mean and error bars standard deviation of five biological replicates composed of three insects, and significant differences between group were assessed by t-test (**** *p*<0.0001; n.s. non-significant).

Previous studies have demonstrated an active involvement of XPO1 during virus infection in vertebrates, invertebrates, and plants, either as a proviral or antiviral factor (Zhang et al., 2021, Yasunaga et al., 2014, Elton et al., 2001, Mathew and Ghildyal, 2017). Specifically, MMV replicates in the nucleus of infected cells in plant and insect hosts and how the nucleocytoplasmic traffic of viral message RNA (vmRNA) and vRNP occurs is still unknown. In order to obtain clues as to whether XPO1 may play a role during MMV infection in *P. maidis*, we first evaluated *PmXPO1* expression in MMV-infected adults across two time points. MMV-infected insects were obtained through microinjection of purified MMV virions in newly emerged adults, males and females. A significant increase in virus abundance was observed from 6 to 12 days after MMV virion microinjection, with a 8.2- and 6.7-fold higher abundance in males and females across time points, respectively, confirming virus replication (**Figure 2c**). Moreover, virus abundance was significantly higher in males than females in both time points analyzed (**Figure 2c**), as previously demonstrated when the virus was acquired through feeding on MMV-infected maize plants (Barandoc-Alviar et al., 2016). However, analysis of *PmXPO1* expression at 6 and 12 days after MMV microinjection demonstrated no significant difference in transcript abundance in both males and females compared to buffer-injected controls (**Figure 2d**). Similarly, a previous study evaluating the global transcriptional response of *P. maidis* to MMV infection, which used high throughput sequencing, did not observe any *PmXPO1* modulation (Martin et al., 2017). These results suggest that if XPO1 plays any function during MMV infection, the basal level of expression would support its role regardless of virus abundance. Furthermore, these results further validate the use of MMV virion microinjection for functional studies which present congruent results compared to those obtained in previous studies using natural virus acquisition (Yao et al., 2019, Martin et al., 2017, Barandoc-Alviar et al., 2016).

### Knockdown of XPO1 reduced *P. maidis* survival in a sex-specific manner

XPO1 plays a vital regulatory role in cell development and physiology through exporting hundreds of cargos from nucleus to cytoplasm, and alterations in XPO1 levels can significantly affect several cellular processes. We further investigated the physiological functions of XPO1 in *P. maidis* through gene silencing using an *in vivo* RNAi approach. First, we tested whether XPO1 knockdown could be efficiently achieved by double-strand RNA (dsRNA) microinjection in whole bodies of adult males and females at transcript and protein levels (**Figure 3a, b**). Upon dsRNA microinjection, the abundance of *PmXPO1* transcripts were significantly reduced by 98% and 87% at 12 days after dsRNA targeting XPO1 (dsXPO1) microinjection in whole bodies of males and females, respectively, compared to dsRNA targeting green fluorescent protein (dsGFP) injected controls (**Figure 3a**). Immunoblot analysis using an anti-XPO1 antibody further confirmed *PmXPO1* silencing at the protein level (**Figure 3b**). Reduction in protein amount was detectable as early as four days after dsRNA microinjection with a more pronounced effect observed at 8 and 12 days after dsXPO1 microinjection, in both males and females whole bodies, compared to control dsGFP-injected insects (**Figure 3b**). These results indicate that *PmXPO1* expression was effectively knocked down at the transcript and protein level in insect bodies.

**Figure 3.**
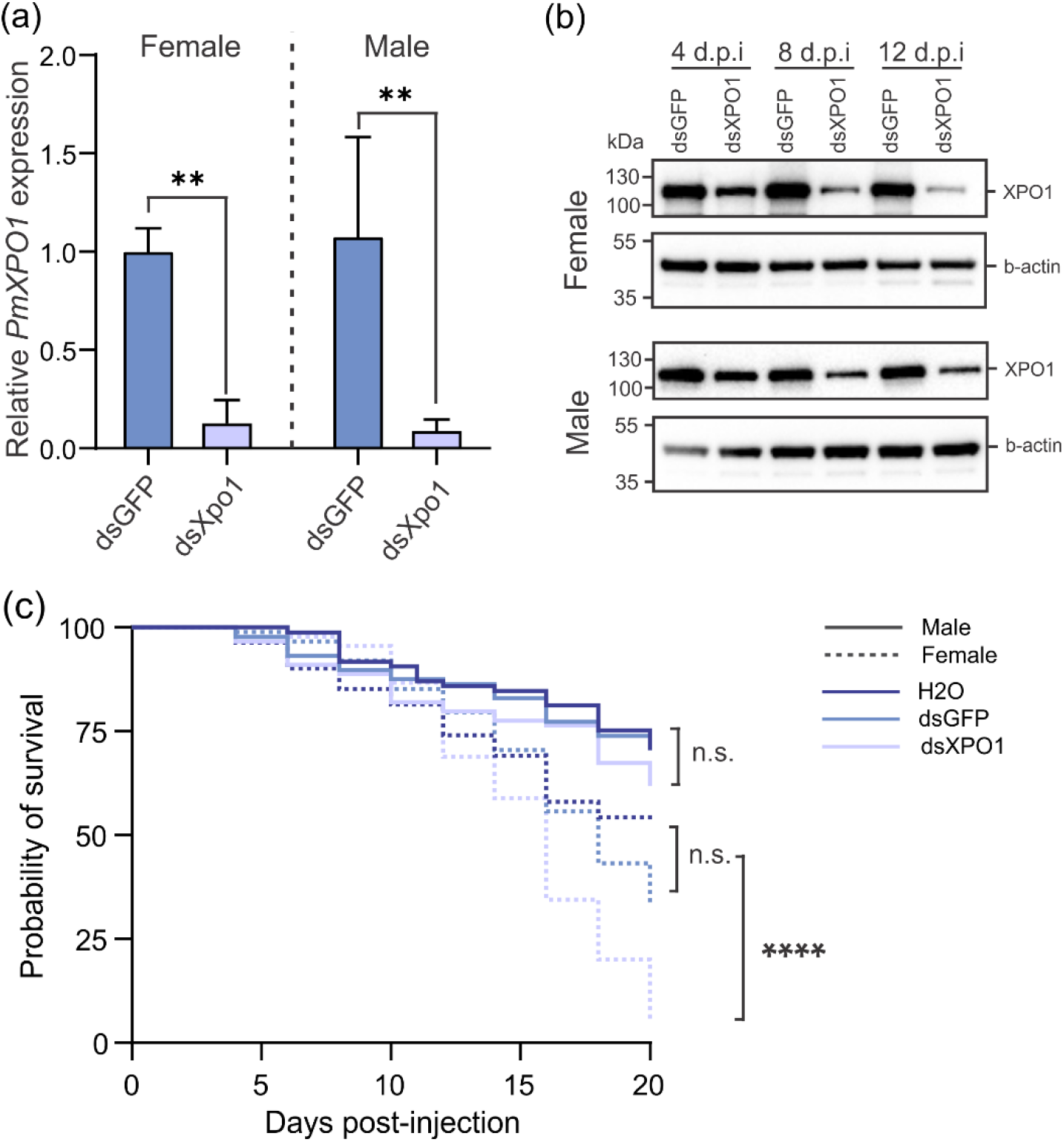
Knockdown of *P. maidis exportin 1* (*PmXPO1*) reduced survival of females but not males. (**a**) Determination of *PmXPO1* knockdown in males and females at 12 days after dsRNA microinjection. Bars indicate mean and error bars standard deviation of three biological replicates composed by three insects. Relative *PmXPO1* expression was quantified by using 2^−ΔΔCT^ and *P. maidis ribosomal protein L10* (*PmRPL10*) used as an internal reference gene. Significant differences between groups were assessed by t-test (** *p*≤0.01). (**b**) Immunoblot analyses of PmXPO1 knockdown evaluated at 4, 8 and 12 days post-injection (d.p.i.) using anti-XPO1 antibody. For each time point a pool of five insects were analyzed. Beta-actin was used as a loading control. (**c**) Kaplan-Meier survival curve of *P. maidis* males and females with *XPO1* knockdown compared to insects microinjected with water and dsGFP, as negative controls. Experiments were independently repeated three times and the pooled data were used to build the survival curves (n = 90 for each treatment). Statistical significance was measured by using Log-Rank test (**** *p*≤0.0001; n.s. non-significant) performed with the survminer package in R software. *P-values* were adjusted using Benjamini & Hochberg (1995) correction for multiple comparisons.

Next, to test whether XPO1-knockdown could have any effect on life span of *P. maidis* male and female individuals, survival of dsRNA-injected newly-emerged adults was determined every other day over a 20-day period (**Figure 3c**). Upon gene knockdown, no significant difference in survival rate was observed for dsXPO1-injected males compared to controls, with >65% survival rate at 20 days after microinjection for all treatments (**Figure 3c**). In contrast, XPO1 knockdown significantly affected female survival compared to water- and dsGFP-injected controls (**Figure 3c**). Survival of dsXPO1-treated females was lower than 6%, whereas approximately 50% of females in control groups (water- or dsGFP-treated) were alive at 20 days after microinjection (**Figure 3c**). These results suggest that XPO1 may play a sex-specific role in the physiology of *P. maidis* significantly affecting survival of female planthoppers.

It was previously demonstrated that pharmacological and genetic XPO1 inhibition extended the lifespan of *Caenorhabditis elegans* by modulating nucleolar dynamics and autophagy (Kumar et al., 2022, Silvestrini et al., 2018). In contrast, our findings demonstrate a sex-specific effect of XPO1 silencing in *P. maidis* physiology negatively affecting female survival, but not males (**Figure 3c**). Whether it is a direct or indirect effect needs to be further investigated. The finding that XPO1 knockdown significantly affected multiple physiological functions in *P. maidis*, including lipid metabolism and reproduction (**Figures 4; 5**), with no apparent effect observed in males, makes it difficult to assign a specific role of XPO1 silencing in modulating *P. maidis* lifespan. Although pharmacological XPO1 inhibition specifically led to increased lifespan of *Drosophila* Sod1 females, a model of amyotrophic lateral sclerosis (ALS; Silvestrini et al., 2018), a direct and conserved role in promoting insect lifespan cannot be generalized. Because Sod1 flies have a very short lifespan compared to wild type individuals (Şahin et al., 2017) and a direct involvement of XPO1 in ALS has been demonstrated (Steyaert et al., 2018, Zhong et al., 2017, Şahin et al., 2017), the effect of XPO1 in fly lifespan may be confounded due to the pleiotropic roles of XPO1 in cell physiology, including ALS. For example, in a *C. elegans* Sod1 model, disruption of the XPO1-mediated nuclear export of Sod1 mutants resulted in higher toxicity in cells and animals (Zhong et al., 2017). Thus, further analysis of wild type flies should be performed in order to clarify the effect of XPO1 in insect lifespan. Of note, the sex specific effect observed here and by Silvestrini et al., 2018, although in opposite directions, is intriguing. It was demonstrated that XPO1 plays an antagonistic pleiotropic role in *C. elegans*, as its inhibition during entire nematode life cycle led to increased mortality (Silvestrini et al., 2018, Büssing et al., 2010). Therefore, the pleiotropic roles of XPO1 in cell physiology may vary according to age across different organisms and further studies are needed to address this topic in insects, including *P. maidis*. Alternatively, XPO1 may play distinct roles in survival across different species. In summary, our study identified a potential target gene to be modulated to control *P. maidis* populations and virus transmission.

**Figure 4.**
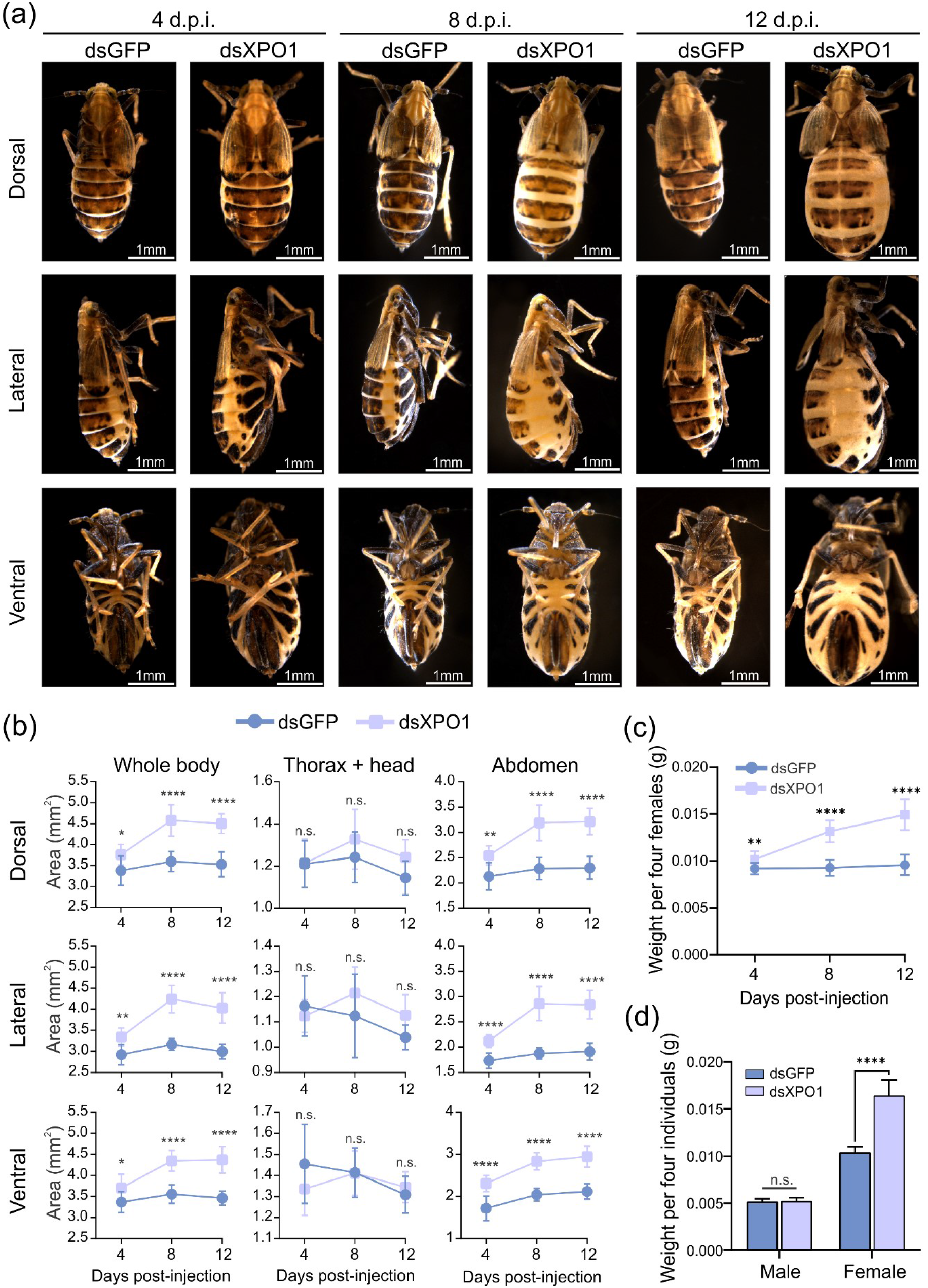
Knockdown of *P. maidis exportin 1* (*PmXPO1*) increased female body size. (**a**) Females with *PmXPO1* knockdown compared to control insects. Representative pictures of females injected with dsGFP or dsXPO1 at 4, 8 and 12 days post-injection (d.p.i.) are shown. (**b**) Body area at different positions (dorsal, lateral, and ventral) for each insect was measured using ImageJ at 4, 8 and 12 days after dsRNA microinjection. Each bar represents mean and standard deviation from nine independent insects. (**c**) Females with *PmXPO1* knockdown were heavier compared to controls. Groups of four insects were weighed at 4, 8 and 12 days after dsRNA injection, as indicated in the figure. Each dot represents the mean and standard deviation of nine to 12 replicates composed by four insects. (**d**) *PmXPO1* knockdown affects females body weight but not males. Insects were weighted 12-days after dsRNA injection. Bars represent mean and standard deviation from 10 replicates composed by four insects each. In **b**, **c** and **d**, significant differences between groups were assessed by two-way ANOVA followed by Sidak’s test (* *p*≤0.05; ** *p*≤0.01; *** *p*≤0.001; **** *p*≤0.0001; n.s. non-significant). One representative experiment is shown from at least two independent experimental replicates for each time point.

### Severe obesity observed in *PmXPO1*-knockdown females

Female dsXPO1-treated planthoppers were clearly larger compared to dsGFP-injected control insects. In order to systematically investigate the effect of XPO1 knockdown on female body size, a time course experiment was performed where the body area and weight was measured at 4, 8 and 12 days after dsRNA microinjection. Adult females with *PmXPO1* knockdown had larger abdomens and were significantly heavier compared to controls at all time points evaluated (**Figure 4**). Significant increases in the body size of females started as early as 4 days after dsXPO1 microinjection compared to dsGFP-injected controls (**Figure 4a, b**). Analysis of the body area of three distinct positions (dorsal, lateral, and ventral) demonstrated a significant increase in size of the whole body and abdomen with no significant difference observed for thorax/head (**Figure 4b**). Furthermore, severe obesity was observed for dsXPO1-treated females with a significant increase in weight starting 4 days after dsXPO1 microinjection, compared to controls (**Figure 4c**). To further verify whether increased weight was a female-specific or general phenomenon observed for both sexes, we evaluated the effect of XPO1 knockdown in male weight at 12 days after dsRNA-microinjection. While dsXPO1-treated females were approximately 55% heavier compared to controls, no significant effect on weight of males was observed (**Figure 4d**). Consistent with increase in weight, glyceride content was specifically and significantly increased in *PmXPO1* knockdown females, but not in males (**Supp. Figure S1**).

Shi et al., 2019 demonstrated that dsRNA-mediated knockdown of *N. lugens* sphingomyelin synthase-like protein 1, which is involved in sphingolipids metabolism, led to increased weight of females, but not of males, suggesting a general sex specific effect of lipid metabolism modulation in planthoppers. In *C. elegans*, XPO1 silencing resulted in elevated lipid accumulation and increased expression of lipases (Silvestrini et al., 2018). Our results demonstrated a higher level of glycerol in female whole bodies and ovaries at 12 days after XPO1 silencing (**Supp. Figure S1**), which may indicate a higher lipase activity. Indeed, a lower level of triglycerides was detected in fat bodies of XPO1-injected females compared to dsGFP-treated control at 12 days after microinjection and further indicates a higher lipase activity. In summary, these results further reinforce the sex-specific effect of XPO1 knockdown on female planthopper physiology, indicating that XPO1 knockdown may either directly or indirectly modulate lipid metabolism.

### *PmXPO1* knockdown negatively affected female reproduction

Impaired lipid metabolism has been correlated with increased abdomen size and reduced fertility in *N. lugens* (Wang et al., 2021, Lu et al., 2018). Because *PmXPO1* knockdown led to abnormal abdomen growth and compromised lipid metabolism in females (**Figures 4; 5**), we further investigated the effect of XPO1 silencing in ovary morphology and reproduction of *P. maidis*. We performed a time course experiment where ovaries were isolated at 4, 8 and 12 days after dsRNA microinjection. Females used in this experiment were approximately three days post-adult eclosion and ovaries isolated from uninjected females immediately before microinjection showed normal development (**Figure 5a**). Similarly to XPO1 silencing in whole bodies, significant reduction in XPO1 transcripts was observed in ovaries of dsXPO1-treated females at 12 days after microinjection compared to control insects (**Figure 5b**). XPO1 silencing resulted in rudimentary ovaries with significant developmental inhibition compared to controls in all time points analyzed (**Figure 5c-h**). Four days after dsRNA microinjection, dsXPO1-treated females exhibited poorly developed ovarioles with no mature eggs or oocytes being observed (**Figure 5c-e**). In contrast, ovaries were fully developed in dsGFP-injected controls, showing healthy ovarioles with typical banana-shaped oocytes (**Figure 5f-h**). No clear difference was observed in external reproductive organs of dsXPO1-injected females compared to control at any of the time points analyzed (**Figure 5i**). These results indicate that XPO1 is required for ovary functions, including oogenesis. Consistent with a major role of XPO1 in ovary functions, oviposition and egg hatching in plants was dramatically reduced in dsXPO1 treated insects compared with control in all time points tested (**Figure 5j, k**). These results further reinforce the role of XPO1 in *P. maidis* reproduction.

**Figure 5.**
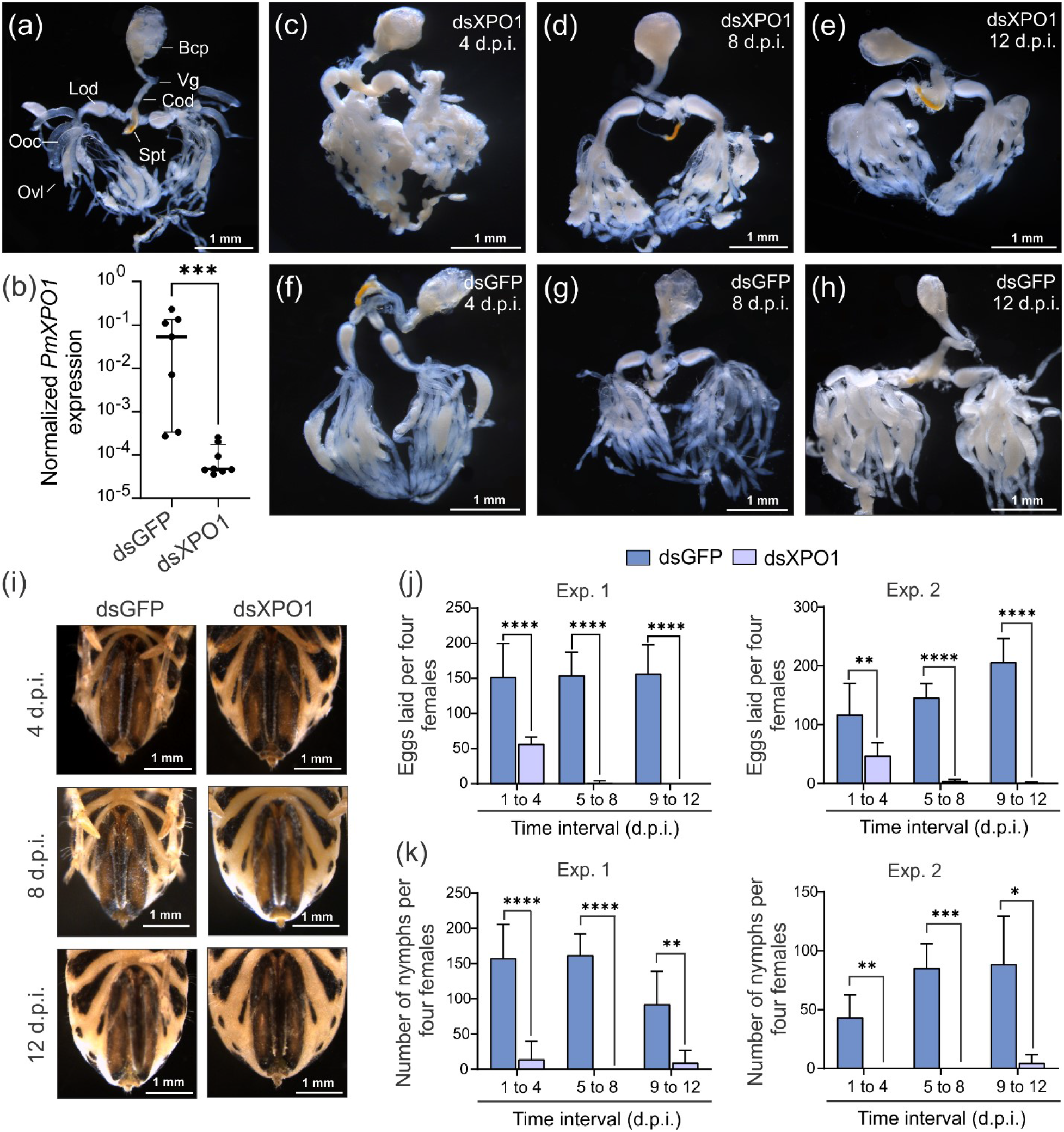
Knockdown of *P. maidis exportin 1* (*PmXPO1*) negatively affected female reproduction. Dissected ovaries are shown for a noninjected individual collected immediately before microinjection (**a**) and for individuals injected with dsXPO1 (**c**-**e**) and dsGFP (**f**-**h**) at 4, 8 and 12 days post-injection (d.p.i.). One representative picture is shown from five dissected ovaries for each treatment from one representative experiment of at least two independent experimental replications for each time point. (**b**) *PmXPO1* expression level was quantified in single dissected ovaries from insects injected with dsGFP or dsXPO1 at 12 d.p.i. Each dot represents a single insect ovary. Error bars represent median and interquartile range. Normalized *PmXPO1* expression was quantified by using 2^−ΔCT^ and *P. maidis ribosomal protein L10* (*PmRPL10*) as an internal reference gene. The nonparametric Mann–Whitney-test was used to test the difference between treatments (*** *p*≤0.001). (**j**) Number of eggs laid at different time intervals after dsRNA injection. Four females were reared in clip cages for oviposition in the second and fourth oldest leaves of two weeks old corn plants and left to lay eggs at time interval of 4 days as indicated in the figure. (**i**) The number of nymphs were quantified after different time intervals after dsRNA microinjection. Four females were released on single corn plants and left to lay eggs at time interval of 4 days as indicated in the figure. Each bar represents the mean and standard deviation of four replicates, each one composed of four insects. Significant differences between groups were assessed by t-test (* *p*≤0.05; ** *p*≤0.01; **** *p*≤0.0001). Bcp: bursa copulatrix; Vg: vagina; Cod: common oviduct; Lod: lateral oviduct, Ovl: ovariole; Ooc: oocyte; Spt: spermatheca.

While a direct role of XPO1 in regulating ovary functions including oogenesis has not been well established in insects reproduction, similarly to our findings, RNAi-mediated XPO1 depletion resulted in significant ovary malformation in *Drosophila* (ElMaghraby et al., 2021). Moreover, Collier et al., 2000 demonstrated that XPO1 is required for *Drosophila* larval progression. Of note, it was suggested that XPO1 could play a vital role in embryogenesis and that the development from embryo to larval stages was likely dependent on maternally inherited XPO1 mRNA from early embryogenesis (Collier et al., 2000). The results obtained here suggest that XPO1 may play a role even earlier in the reproduction cycle of *P. maidis* by directly regulating oogenesis rather than overall ovary development. Whereas no clear difference was observed in the morphology of most ovary parts, including bursa copulatrix, vagina, common and lateral oviduct and spermatheca, ovariole development and oocytes were drastically compromised upon dsXPO1 microinjection (**Figure 5c-h**). Nevertheless, because our experiments were performed in newly-emerged females, with well-developed ovaries (**Figure 5a**), silencing of XPO1 in nymphal stages of *P. maidis* should be performed in order to verify its effect in ovary morphogenesis. XPO1 plays multiple conserved roles in cell physiology including regulation of cell cycle progression, ribosomal synthesis, and transcription. Furthermore, Kırlı et al., 2015 identified a tremendously large number of XPO1 cargo in *Xenopus oocytes* uncovering putative new functions including autophagy, peroxisome biogenesis, cytoskeleton, ribosome maturation and mRNA degradation. Moreover, the involvement of XPO1 in the biogenesis of the nuclear pore complex has been demonstrated during *Drosophila* oogenesis (Hampoelz et al., 2019). Therefore, it is likely that XPO1 knockdown would affect basic processes involved in reproduction, such as oogenesis.

Other cellular pathways relying on XPO1 regulation through its nuclear export activity have been characterized as controlling basic cellular functions, including reproduction. For example, XPO1 mediates the nuclear export of unprocessed PIWI-interacting RNA (piRNA) precursors by interacting with nuclear RNA export factor 3 (Nxf3) adapter protein, and *nxf3*-null mutant flies were sterile (ElMaghraby et al., 2019). Therefore, interference with XPO1 activity could affect piRNA biogenesis and consequently insect reproduction. Furthermore, XPO1 is involved in micro-RNA (miRNA) processing in *C. elegans*, *Drosophila* and human cells (ElMaghraby et al., 2019, Martinez et al., 2017, Büssing et al., 2010). Pharmacological and genetic XPO1 inhibition caused vulval bursting and sterility by affecting primary-miRNA-to-pre-miRNA processing in *C. elegans* (Büssing et al., 2010). In *N. lugens*, silencing of core components of miRNA pathway drastically affected insect development and reproduction (Wang et al., 2023a). RNAi-mediated silencing of *Argonaute1*, *Dicer1* and *Drosha* caused impaired ovary and oocytes development with significant reduction of laid eggs (Wang et al., 2023a). Similarly, *Dicer1* silencing negatively affected ovary development and fecundity of the white-backed planthopper, *Sogatella furcifera* (Zeng et al., 2023). Interestingly, phenotypes observed in *N. lugens* ovaries looked similar to the ones observed in *P. maidis* suggesting that a similar pathway could be modulated through XPO1 knockdown. Finally, lipid storage is critical for insect reproduction and impaired lipid metabolism has been demonstrated to directly and indirectly affect *N. lugens* reproduction (Wang et al., 2021, Lu et al., 2018). Our findings demonstrate a significant modulation of lipid metabolism with females becoming obese over time and this could partially explain impaired *P. maidis* reproduction. Although there are several possible explanations for the effect of XPO1 on *P. maidis* reproduction, they are not mutually exclusive, and the combination of different mechanisms may be a more likely explanation considering the pleiotropic roles of XPO1 in cell physiology. Further investigation is needed to reveal the mechanism by which XPO1 affects *P. maidis* reproduction.

### *PmXPO1* knockdown reduced MMV accumulation in insect whole body

Although *PmXPO1* was not modulated during MMV infection in the whole body of male and female individuals (**Figure 2d**), we further investigated whether XPO1 plays a role in MMV infection by gene silencing followed by analysis of MMV accumulation. If MMV co-opts *PmXPO1*-encoded protein to mediate nuclear exporting of viral proteins or vRNPs, silencing of *PmXPO1* would negatively affect virus abundance. In order to address this hypothesis, newly emerged adults, males and females, were simultaneously microinjected with a mix containing dsXPO1 and MMV virions, and gene silencing and MMV accumulation were analysed 12 days after microinjection through RT-qPCR (**Figure 6**). As previously shown for microinjection of dsXPO1 alone (**Figure 3a, b**), upon dsXPO1 and MMV virions microinjection, efficient XPO1 silencing was observed in whole body of males and females with higher than 90% reduction of XPO1 transcripts, at 12 days after microinjection (**Figure 6a**). Analysis of virus abundance based on RT-qPCR demonstrated a significantly lower MMV abundance in dsXPO1-treated males, with approximately 50% reduction compared with dsGFP-injected control (**Figure 6b**). However, no significant effect was observed on MMV abundance in female whole-bodies (**Figure 6b**). These results were consistently observed across four independents experimental replicates. Because MMV accumulates in lower levels in females compared to males (**Figure 2c**; Barandoc-Alviar et al., 2016) and residual amount of PmXPO1 could sustain virus infection, we sought to further investigate the effect of *PmXPO1* knockdown in MMV accumulation at earlier and later time points during virus infection. A time course experiment was performed and MMV abundance was evaluated at 6, 12 and 15 days after dsRNA plus MMV virion microinjection. Significant reduction in XPO1 transcripts was observed at all time points analysed for both sexes (**Figure 6c, d**). Whereas no significant effect in virus abundance was observed at 6 days after microinjection for both sexes (**Figure 6e, f**), a significant reduction in MMV abundance was observed at 12 and 15 days in males (**Figure 6e**), consistent with the previous results (**Figure 6b**). Interestingly, no significant effect in MMV abundance was observed in females at 12 days after microinjection, whereas significantly lower virus abundance was detected at 15 days after microinjection (**Figure 6f**). These results suggest that XPO1 plays a proviral role during MMV infection in its insect vector.

**Figure 6.**
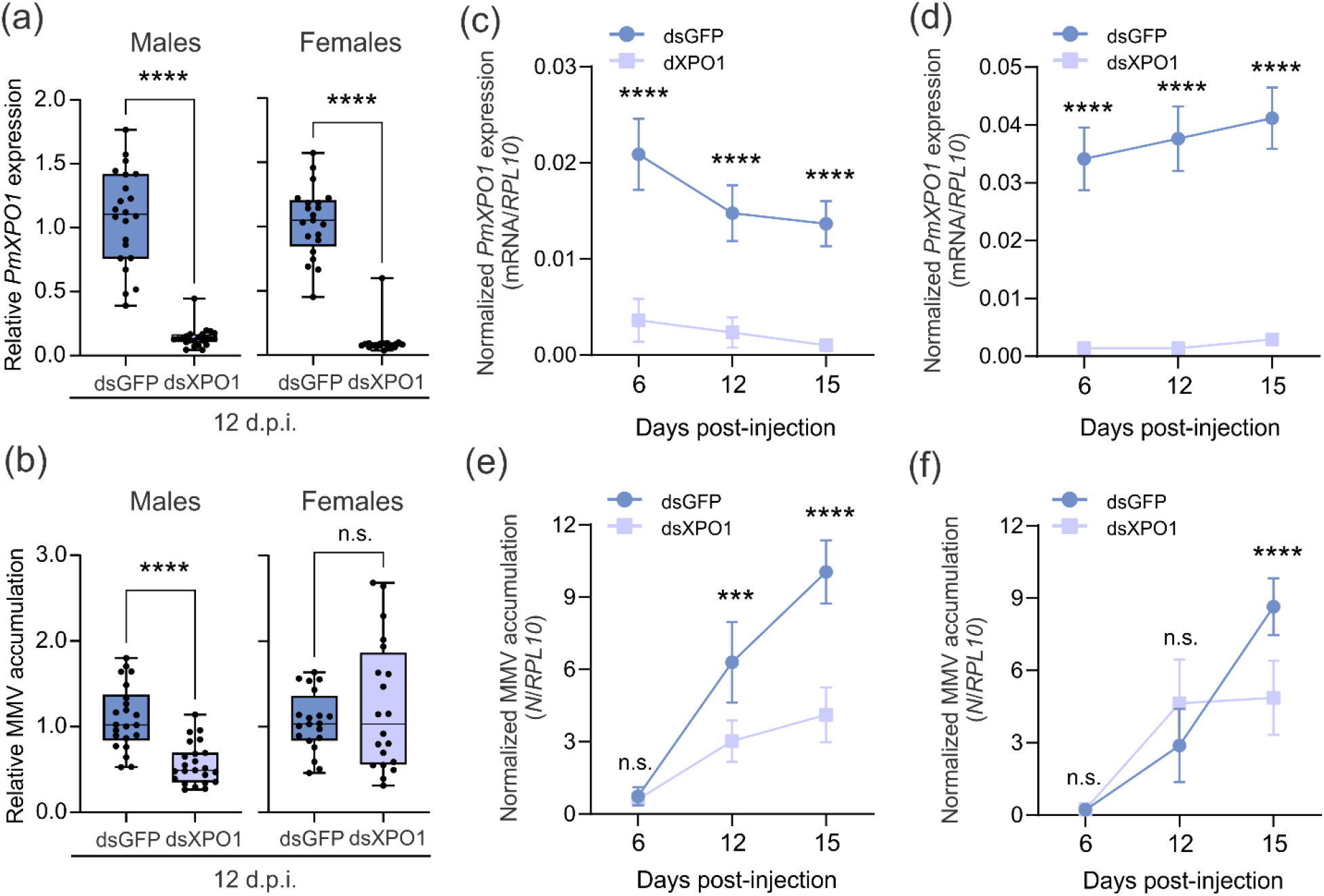
Knockdown of *P. maidis exportin 1* (*PmXPO1*) negatively affected maize mosaic virus (MMV) accumulation. (**a**) Determination of relative transcripts level of *PmXPO1* and (**b**) MMV accumulation at 12 days after dsRNA and MMV virions microinjection, for males and females. Each dot represents a biological replicate composed of three insects. Boxes represent 25th to 75th percentiles, horizontal bar median, and whiskers the minimum and maximum values. Relative *PmXPO1* expression and MMV accumulation were quantified by using 2^−ΔΔCT^ and *P. maidis ribosomal protein L10* (*PmRPL10*) used as an internal reference gene. Significant differences between groups were accessed by t-test (**** *p*≤0.0001; n.s. non-significant). d.p.i.: days post-injection. (**c**-**f**) Time course experiment determining *PmXPO1* expression (**c** and **d**) and MMV accumulation (**e** and **f**) at 6, 12 and 15 days after dsRNA and MMV virion microinjection in males (**c** and **e**) and females (**d** and **f**) adults. Circles and squares represent mean and error bars standard deviation of five to seven biological replicates composed by three insects. Insects were injected with 80 nl of 1000 ng/µl of dsRNA plus 100ng/µl of MMV virions. Normalized *PmXPO1* expression and MMV accumulation were quantified by using 2^−ΔCT^ and *PmRPL10* used as an internal reference gene. Significant differences between groups were assessed by two-way ANOVA followed by Sidak’s test (*** *p*≤0.001; **** *p*≤0.0001; n.s. non-significant).

MMV replicates in the nucleus of infected cells in insect and plant hosts and the mechanism by which vRNP, viral proteins e vmRNA traffic from nucleus to cytoplasm is still unclear. In contrast with the general antiviral response against arbovirus infection (Yasunaga et al., 2014), our results suggested that XPO1 has proviral activity during MMV infection in *P. maidis* as its knockdown led to lower virus accumulation. Previous studies have demonstrated that viruses co-opt XPO1 for distinct purposes and in distinct stages during the viral life cycle. XPO1 has been demonstrated to mediate nucleocytoplasmic export of vRNP. For example, for influenza virus, a RNA virus that replicates in the nucleus of infected cells, the nuclear export of vRNP has been shown to be mediated by XPO1 interaction with the viral nuclear export protein (NEP) and required for cytoplasmic virion maturation at late stages of virus infection (Huang et al., 2013, Elton et al., 2001). Moreover, the interaction of XPO1 with vRNP of HBV, a DNA virus whose replication takes place in the nucleus-cytoplasm, is required in early and late events of virus infection (Yang et al., 2022, Su et al., 2022). Although localization of MMV proteins is conserved in plant and insect cells (Martin and Whitfield 2018), how nucleocytoplasmic traffic occurs is poorly understood in both hosts. In plants, vRNPs are speculated to be the core units exiting the nucleus (Jackson et al., 2005, Zhou et al., 2019), and the interaction of the PYDV matrix protein with XPO1 reinforces a putative role of XPO1 in mediating vRNP nuclear export (Anderson et al., 2018). In contrast, it has been hypothesized that MMV buds out of the perinuclear space to traffic through the cytoplasm to reach cell periphery as vRNP in insect cells, which would occur independently of XPO1 (Jackson et al., 2005). Alternatively, MMV vRNP may rely on XPO1 to exit the nucleus through NPC in order to spread to adjacent cells, which must be further investigated.

Furthermore, XPO1-mediated export of viral transcripts via adapter proteins has been demonstrated. The export of HIV unspliced viral transcripts is mediated by XPO1 through its interaction with the viral adapter Rev protein (Yang et al., 2022, Su et al., 2022, Neville et al., 1997, Fischer et al., 1995). MMV vmRNA are structurally similar to eukaryotic mRNA, consisting of a methylated guanosine (m^7^G) cap and poly-A tail at 5’ and 3’ end, respectively. Thus it is likely that MMV hijacks the canonical mRNA export (e.g. mediated by Nxt1/Nxtf1) host machinery instead of relying on a non-canonical mRNA export pathway, for example XPO1-mediated. Lastly, XPO1 regulates the nucleocytoplasmic distribution of several viral proteins, for example the matrix protein of RSV (Ghildyal et al., 2009), NS5 protein of dengue virus (Pryor et al., 2006) and phosphoprotein of rabies virus (Pasdeloup et al., 2005) have been demonstrated to be regulated by XPO1 export activity. Thus, although we provide evidence of direct involvement of XPO1 in MMV infection, further investigations will be performed to characterize the mechanism. Previous work demonstrating an XPO1 interaction with animal (Pasdeloup et al., 2005) and plant (Anderson et al., 2018) rhabdoviruses, including matrix and phosphoprotein, respectively, further reinforces the new finding that XPO1 may play a direct role in MMV infection in a cross kingdom manner.

## CONCLUDING REMARKS

XPO1 plays multiple regulatory roles in cell functions and in virus infection, and we demonstrated that RNAi-mediated silencing of *XPO1* simultaneously and negatively affected virus accumulation and insect physiology. This study provides the first evidence that XPO1 plays a proviral role during plant rhabdovirus infection in its insect vector. XPO1 may be a potential dual target gene to simultaneously control arbovirus infection and the insect vector by interfering with insect survival and reproduction. MMV is transmitted in a dose-dependent manner and reducing MMV accumulation in the insect vector has potential to block virus transmission. Further experiments to elucidate the mechanism by which XPO1 positively affects MMV accumulation and impairs insect reproduction are underway. Because of the XPO1 conserved role mediating nuclear export in eukaryotic cells, we speculate that XPO1 may play a cross kingdom role in modulating MMV infection in plants and insects cells.

## EXPERIMENTAL PROCEDURES

### Insect colony and virus

The corn planthopper colony used in this study was originally established from a field collected colony at University of Hawaii, Honolulu (Barandoc-Alviar et al., 2016, Namba and Higa, 1971). Insects were reared on healthy 3- to 4-week-old maize plants (cv. Early Sunglow) and maintained in insect-proof cages under a 12:12 light/dark photoperiod at 26°C in a reach-in growth chamber (Conviron), as previously described (Yao et al., 2019). To synchronize insect age, mature adults were placed on healthy 3- to 4-week-old maize plants for a 3 day-oviposition period, and then removed from the plants and eggs let to hatch. For all experiments, short-winged insects approximately 1- to 3-days after becoming adults from the age calibrated colony were used unless stated otherwise. All experiments were conducted under the same environmental conditions (photoperiod and temperature) described above for insect colony maintenance.

Maize mosaic virus (MMV; GenBank accession number MK828539) virions were obtained from MMV-infected maize leaves (cv. Early Sunglow) through a combination of differential and sucrose density gradient centrifugations, as previously described (Yao et al., 2019, Ammar et al., 2005). Virions were stored in a ultra-low temperature freezer at −80°C and thawed on ice before microinjection procedures.

### Identification and molecular characterization of *PmXPO1*

Identification of *PmXPO1* coding sequence was performed through a local BLASTn search in two previously assembled *P. maidis* transcriptomes (Wang et al., 2023b, Martin et al., 2017). The nucleotide sequence of *D. melanogaster exportin 1* (*DmXPO1*, access number NM_001316387.1) was used as a query. Potential *P. maidis* transcripts showing significant similarity (*E* < 10^−3^) to *DmXPO1* were further analyzed for the presence of potential open reading frames (ORFs) with Translate tool implemented in Expert Protein Analysis System (ExPASy; Gasteiger et al., 2003) and compared against the NCBI non-redundant protein sequence database using BLASTx. Identification and domain annotation was performed with the Simple Modular Architecture Research Tool (SMART; Letunic et al., 2020) and compared to sequences of other previously characterized XPO1 sequences from vertebrates and invertebrates. Pairwise identity comparisons of full-length amino acid sequences of XPO1 were performed with the program Species Demarcation Tools (SDT v. 1.2; Muhire et al., 2014) using the MUSCLE alignment option.

To validate the *in silico* identified *PmXPO1*, the predicted ORF was amplified using reverse transcriptase-polymerase chain reaction (RT-PCR) with specific primers designed according to the obtained transcripts sequences (**Supp. Table 1**). PCR was performed using complementary DNA (cDNA) template synthesized from 1 µg of total RNA extracted from a mix of males and females insects, as described below. The final reaction consisted of 1 µl of undiluted cDNA, 0.25 µM of forward and reverse primers mixed in a final volume of 25 µl reaction using Q5 Hot Start High-Fidelity 2X Master Mix (NEB), following the manufacturer’s instructions. PCR products were checked on 1% agarose gel and directionally cloned using pENTR/D-TOPO Cloning Kit (Invitrogen) and then transformed into 5-alpha competent *Escherichia coli* cells (NEB), according to the manufacturer’s instructions. Positive clones were screened through colony PCR using gene specific primers (**Supp. Table S1**) and bidirectionally sequenced by the Genomic Sequencing Laboratory at North Carolina State University (GSL-NCSU).

### Phylogenetic analysis

To investigate the evolutionary relationship of the identified sequence, a phylogenetic analysis was performed. The final data set included insect sequences closely related to *PmXPO1*, which were selected based on BLASTp analysis, and representative sequences from the orders Blattodea, Diptera, Hemiptera, Hymenoptera, Orthoptera, and Thysanoptera. Additionally, to verify the level of divergence between insects XPO1 and those from more distantly related organisms, two sequences from vertebrates and three sequences from plants were also added to the analyzes. Representative sequences of other exportins from karyopherin-β subfamily (exportin-5 and exportin-T) closest related to XPO1 were added into the phylogeny as an outgroup (O’Reilly et al., 2011). For further sequence information used in the phylogenetic analysis, see **Supp. Table S2**.

Phylogenetic analysis was conducted according to Xavier et al., 2021. Briefly, multiple alignment of full-length XPO1 amino acid sequences was performed using the MUSCLE algorithm implemented in MEGA X (Kumar et al., 2018). Phylogenetic trees were constructed with MrBayes 3.2.6 software and estimate of the amino acid model was automatically conducted by setting the prior for the amino acid model to mixed, as implemented in MrBayes (Ronquist et al., 2012). The analysis was carried out running two independent runs of 10,000,000 generations with sampling at every 1,000 generations and a burn-in of 25%. Convergence between runs were accepted when the average standard deviations of split frequencies was lower than 0.01. The phylogenetic tree was visualized and edited using FigTree and Corel Draw, respectively.

### RNA extraction, cDNA synthesis and RT-qPCR

For total RNA extraction, pooled insects whole bodies or tissues were homogenized using the Tissue Lyser II (Qiagen) in a 1.7 ml microcentrifuge tube containing 500 µl of ice-cold trizol and five three-millimeter glass beads (Pyrex). Samples were homogenized at room temperature for two minutes at oscillation frequency of 30 hertz. Total RNA was extracted using Trizol Reagent (ThermoFisher) followed by ethanol precipitation, according to manufacturer’s instructions. Total RNA samples were treated with DNase-I using TURBO DNA-free Kit (Invitrogen), according to manufacturer’s instructions. The concentration and purity of total RNA was measured with a NanoDrop One^©^ spectrophotometer (ThermoFisher), and the concentration adjusted to 100 ng/µl with nuclease-free water. One microgram of total RNA was used for the first strand cDNA synthesis using the Verso cDNA Synthesis Kit (ThermoFisher) with RT-enhancer to remove residual genomic DNA, following the manufacturer’s instructions. The cDNA was diluted 8-fold in nuclease-free water prior to quantification by RT-qPCR. The RT-qPCR reactions consisted of 4 μl of diluted cDNA, 1 μl of mixed primers containing 5 μM of each primer (forward and reverse) and 5 µl of iTaq Universal SYBR Green Supermix (BioRad) in a final volume of 10 μl. The RT-qPCR was carried out with a CFX Connect Real-Time System (BioRad), and cycles consisted of an initial denaturation step at 95°C for 1 min, followed by 40 cycles of 95°C for 15 sec and 60°C for 1 min. For all reactions, amplification specificity was checked based on melting curve analysis using default settings. All reactions were performed in duplicates, and a coefficient of variation of 1% was used as a cutoff to verify the reliability of the technical replications. *PmXPO1* expression levels and virus abundance were quantified by either the comparative cycle threshold method (2^−ΔΔCT^) or normalized RNA abundance (2^−ΔCT^), using the *P. maidis ribosomal protein L10* (*PmRPL10*) as an internal reference gene (Livak and Schmittgen, 2001, Barandoc-Alviar et al., 2016). All primers used here were first tested for efficiency using a serial dilution of cDNA (5^1^ to 5^5^) and the quantification cycle values obtained were used to calculate the efficiency based on the standard curve method. Primer pairs with efficiencies between 90% and 110% and showing a single peak based on melting curves, were accepted for our analysis.

### *PmXPO1* expression analyses

To evaluate the *PmXPO1* expression pattern across developmental stages and tissues, samples were collected from nymphs 1st to 5th instars, adult males and females, and female ovaries and guts. According to the developmental stage, a variable number of individuals were pooled to compose a biological replicate (1st stage nymph n = 20, 2nd nymph n = 10, 3rd nymph n = 10, 4th nymph n = 5, 5th nymph n = 5, adult males n = 3 and adult females n = 3). Females were anesthetized on ice and tissues were dissected under a stereomicroscope Leica M205C in ice-cold 60% ethanol followed by two washing steps in 60% ethanol and then five ovaries and six guts were separately pooled to compose a biological replicate. Total RNA was extracted using Trizol Reagent and *PmXPO1* expression was evaluated from five biological replicates by RT-qPCR as described above.

To evaluate the effect of MMV infection on *PmXPO1* expression, MMV-infected males and females were obtained by microinjection of 50 nl of 100 ng/µl of purified MMV virions, as described below. As a control, a group of insects were injected with 50 nl of buffer. After microinjection, insects were reared in healthy maize plants and collected at six and 12 days after virus microinjection. For each sample, total RNA was extracted from five biological replicates composed of pools of three insects each using Trizol Reagent and *PmXPO1* expression and virus accumulation were evaluated by RT-qPCR as described above.

### dsRNA synthesis and microinjection

Double-stranded RNA was synthesized with T7 RiboMAX Express RNAi System, according to manufacturer’s protocols. Briefly, sets of primers with T7 RNA polymerase promoter sequence added to the 5’ ends and without were designed to amplify 445 nt in the central region of the *PmXPO1* coding sequence (**Supp. Table 1**). Primers were designed following the T7 RiboMAX Express RNAi System guidelines and combined in two separate PCR reactions, forward-T7/reverse or forward/reverse-T7, to generate the template for *in vitro* transcription. PCR reaction was performed using 1 μl of undiluted cDNA template synthesized from 1 µg of total RNA extracted from a mix of male and female individuals using Q5 Hot Start High-Fidelity 2X Master Mix (NEB), as described above. For each primer combination, three 25 μl PCR reactions were pooled and purified using Monarch PCR & DNA Cleanup Kit (5 μg; NEB), according to manufacturer’s instructions. Specificity and concentration of purified PCR products were checked on 1% agarose gel and with a NanoDrop One^©^ spectrophotometer (ThermoFisher), respectively. Approximately 600 ng of each purified PCR product was used as template for *in vitro* dsRNA synthesis according to the manufacturer’s protocol. Double-stranded RNA was purified using Monarch RNA Cleanup Kit (50 µg; NEB) and eluted in 30 μl of nuclease-free water. Specificity and integrity of dsRNA were checked on 1% agarose gel and quantified with a NanoDrop One^©^ spectrophotometer (ThermoFisher). Double-stranded RNA targeting the gene encoding green fluorescent protein (GFP) was synthesized and used as a control.

Insects microinjections were performed according to Yao et al., 2019, with slight modifications. Briefly, insects were anesthetized on ice and held with a micro tweezer on the microscope stage and then injected with dsRNA and/or MMV virions at speed of 50 nl/sec using a Nanoinjector III (Drummond Scientific) under a Leica S APO stereo microscope. Insects were injected on the membrane between the meso- and meta-thoracic legs and allowed to recover for one day after microinjection on healthy maize plants to reduce death from microinjection-associated damage before experiments were performed. Insects were injected with 80 nanoliter of 1,000 ng/µl of dsXPO1 or dsGFP unless stated otherwise.

### Effect of *PmXPO1* knockdown on MMV accumulation

To evaluate the effect of *PmXPO1* knockdown on MMV accumulation, virus abundance and *PmXPO1* silencing levels were evaluated in males and females whole bodies at 12 days after microinjection. Insects were simultaneously injected with 80 nl of a mixture containing 1,000 ng/µl of dsXPO1 or dsGFP plus 100 ng/µl of purified MMV virions. Total RNA was extracted from pools of three insects using Trizol Reagent and *PmXPO1* expression and virus accumulation were evaluated by RT-qPCR as described above. MMV accumulation was determined using primers to detect the *MMV-N* gene (**Supp. Table S1**). To verify whether *PmXPO1* knockdown affected MMV accumulation at early and later time points after microinjection, a time course experiment was performed and MMV accumulation evaluated at 6, 12 and 15 days after microinjection.

### Effect of *PmXPO1* knockdown on *P. maidis* reproduction and phenotype

To verify the effect of *dsXPO1* knockdown on female reproduction, the number of eggs laid and hatched nymphs were evaluated at time intervals of 1 to 4, 5 to 8 and 9 to 12 days after dsRNA microinjection. Approximately 3 days after becoming adults, individual females were injected with 80 nl of 1,000 ng/µl of dsXPO1 or dsGFP. For oviposition assays, two groups of four females were separately reared on the 2nd and 3rd oldest leaves of a 2-weeks old maize plant (cv. Early sunglow) using a rectangular clip cage (1.75” x 2” x 0.625”) placed closest to the base of the leaf. After each time interval, leaves were collected and stored in 100% ethanol at 4°C until further processing. Eggs were visualized and counted under a stereomicroscope Leica M205C following staining with McBride’s solution and destaining with lactic acid:glycerol:water (1:1:1), as previously described (Backus et al., 1988). Four independent maize plants were used for each treatment, and two leaves from each plant were analyzed. To quantify the number of eggs hatching, four females were transferred onto a single 7-day old maize plant (cv. Early Sunglow) and given an oviposition period according to each time interval. After the oviposition period, insects were then removed and plants allowed to grow, and the number of hatched nymphs counted. Maize plants used in this experiment were grown in a cone-tainer (1.5” x 8.25”) and covered with a plastic tube cage capped with insect-proof mesh. Oviposition and hatching assays were performed under a 12:12 light/dark photoperiod at 26°C in a reach-in plant growth chamber. These experiments were independently repeated at least twice for each time interval.

Insect morphology and weight were analyzed at 4, 8 and 12 days after dsRNA microinjection. Insects were anesthetized on ice and imaged with a Leica MC 190 HD camera coupled to a stereomicroscope Leica M205C. For each insect, three photos were taken in different positions (dorsal, lateral, and ventral). Area of thorax + head, abdomen and whole body of females were measured using ImageJ software (Schneider, C. A. et al., 2012). Groups of four females were weighed at 4, 8 and 12 days after dsRNA microinjection using a precision scale. Male weights were recorded at 12 days after dsRNA microinjection, as a control.

To verify ovary morphology, females were dissected at 4, 8 and 12 days after dsGFP and dsXPO1 microinjection under a stereomicroscope in ice-cold 60% ethanol. Photos were taken immediately after dissection with a Leica MC 190 HD camera coupled to the stereomicroscope.

### Efficiency of *PmXPO1* knockdown in ovaries

To quantify *PmXPO1* knockdown efficiency in ovaries, females were dissected 12 days after dsGFP and dsXPO1 microinjection under a stereomicroscope Leica M205C in ice cold 60% ethanol and washed twice before being homogenized. Single ovaries were thoroughly homogenized with a disposable pestle in a microcentrifuge tube containing 35 µl of DNase/RNase-free distilled water (Invitrogen) followed by addition of 35µl of 50% slurry (weight/volume) of Chelex 100 (Bio-Rad). The homogenate was briefly vortexed and heated for 5 minutes at 94°C and centrifuged at 13,000 *g* at 4°C for five minutes. Ten microliters of the supernatant were used for cDNA synthesis using Verso cDNA synthesis kit (ThermoFisher), following manufacturer’s instructions. Gene expression was quantified through RT-qPCR using four microliters of four-fold diluted cDNA, as described above. This experiment was repeated two independent times.

### Glycerolipid quantification

Glycerolipid content of male and female whole bodies and female ovaries and fat bodies was measured 12 days after dsGFP and dsXPO1 microinjection using the colorimetric Serum Triglyceride Determination kit (Sigma, Triglyceride Reagent T2449; Free Glycerol F6428; Glycerol Standard Solution G7793). Female ovaries and fat bodies were dissected under a stereomicroscope Leica M205C in ice-cold PBS 1X and three ovaries or fat bodies were pooled to compose a biological replicate. Fat bodies were dissected in 100 µl of PBS 1X by removing internal organs, thorax, head, and legs while retaining the abdomen cuticle and adhered fat body tissues. Ovaries were washed twice in PBS 1X before being homogenized. For whole-body males and females, three individuals were pooled to compose a biological replicate.

Pooled tissue samples were homogenized with a disposable pestle in a microcentrifuge tube containing 100 µl of ice-cold PBST (1X PBS + 0.5% Tween 20). The homogenate was incubated at 70°C for 10 minutes and spun down for 30 seconds at 587*g*. Samples were kept on ice during the whole process to avoid enzymatic degradation of glycerolipid into free glycerol by endogenous enzymes.

The quantification of triglycerides using the colorimetric assay was performed according to manufacturer’s instructions with slight modifications. Briefly, three 10 µl replicates of each biological replicate were pipetted into individual wells of a clear-bottom 96-well plate (BRANDplates). Standard curves were prepared using serial dilutions of triolein equivalent glycerol standard (2.0, 1.0, 0.5, 0.25, 0.125, 0.0625 and 0 mg/mL), and three 10 µl replicates of each dilution were pipetted into individual wells of the same 96-well plate, as performed for biological replicates. To quantify the free glycerol, 100 µl of Free Glycerol Reagent were added in each well and mixed followed by five minutes incubation at 37°C. Initial absorbance was recorded at 540 nm using Gen5 plate reader (BioTek). To quantify the total triglycerides, 20 µl of Triglyceride Reagent were sequentially added in each well and mixed followed by incubation at 37°C for five minutes and then final absorbance were recorded as described above. Standard curves were obtained by regression analysis of corrected absorbance at 540 nm values for a given dilution in relation to the concentration of triolein standard in milligrams in each dilution. This experiment was repeated two independent times.

### Immunoblot analysis

To quantify PmXPO1 knockdown efficiency at protein level, immunoblot analysis was performed according to Kanakala et al., 2023. For total protein extraction, three pooled insects were homogenized with a disposable pestle in a microcentrifuge tube containing 320 µl of 2x Laemmli Sample Buffer (BioRad) and 5% of 2-mercaptoethanol. The homogenate was incubated at 100°C for 10 minutes and spun down for 5 minutes at 5,000 rpm in a benchtop microcentrifuge. Ten microliters of the supernatant were loaded onto polyacrylamide gels using TGX FastCast Acrylamide kit, 10% (BioRad) and run for 1 hour at 150 volts. Proteins were transferred to a LPF PVDF membrane using the Trans-Blot Turbo RTA Transfer kit (Bio-Rad), following the manufacturer’s instructions. Membranes were blocked with 5% non-fat dry milk overnight at 4°C under gentle agitation followed by 1,5 hours incubation at room temperature with anti-CRM1 (MA5-27879, Invitrogen, 1:2000 dilution) antibody to detect PmXPO1 protein. After three 10 minutes washing steps with TBS-T, membranes were probed with the alkaline phosphatase-conjugated secondary antibody goat-anti mouse (CRM1, 1:2000 dilution) for 1,5 hours at room temperature. After that, membranes were washed three times with TBS-T and chemiluminescent detection was performed with ProSignal Pico (GeneseScientific) following the manufacturer’s instructions. Images were acquired with iBright Imaging system (CL1000, ThermoFisher). As a loading control, membranes were stripped using Restore Western Blot Stripping Buffer (ThermoScientific) and reprobed with anti-beta actin (MA5-15739-HRP, 1:2000) following the steps described above.

### Statistical analysis

Statistical analyses were performed using GraphPad Prism 9.0.0 software. We used t-test to evaluate single variable under two experimental conditions, one-way ANOVA for single variable across multiple conditions and two-way ANOVA followed by Sidak’s test when two or more variables were being analysed and selected pairs of means were compared, unless stated otherwise. For gene expression and virus accumulation, values were log-transformed prior statistical analysis when needed and non-transformed values were plotted in all cases. A significance level of 0.05 was used.

## ACKNOWLEDGEMENTS

We thank members of the Plant-Virus-Vector Interactions Lab team, especially Dr. Dorith Rotenberg and Dr. Jinlong Han, for the valuable discussion throughout the course of the project. The North Carolina State University, Department of Entomology and Plant Pathology, was part of a team supporting DARPA’s Insect Allies Program. The views expressed are those of the authors and should not be interpreted as representing the official views or policies of the U.S. Government.

## AUTHORS CONTRIBUTION

**Cesar A.D. Xavier:** conceptualization, methodology, investigation, formal analysis, validation, visualization, writing – original draft preparation, writing – review & editing. **Clara Tyson**: methodology and investigation. **Leo M. Kerner:** methodology and investigation. **Anna E. Whitfield**: conceptualization, methodology, investigation, funding acquisition, project administration, supervision, writing – original draft preparation, writing – review & editing

## SUPPLEMENTARY MATERIAL

**Supplementary Table S1.**
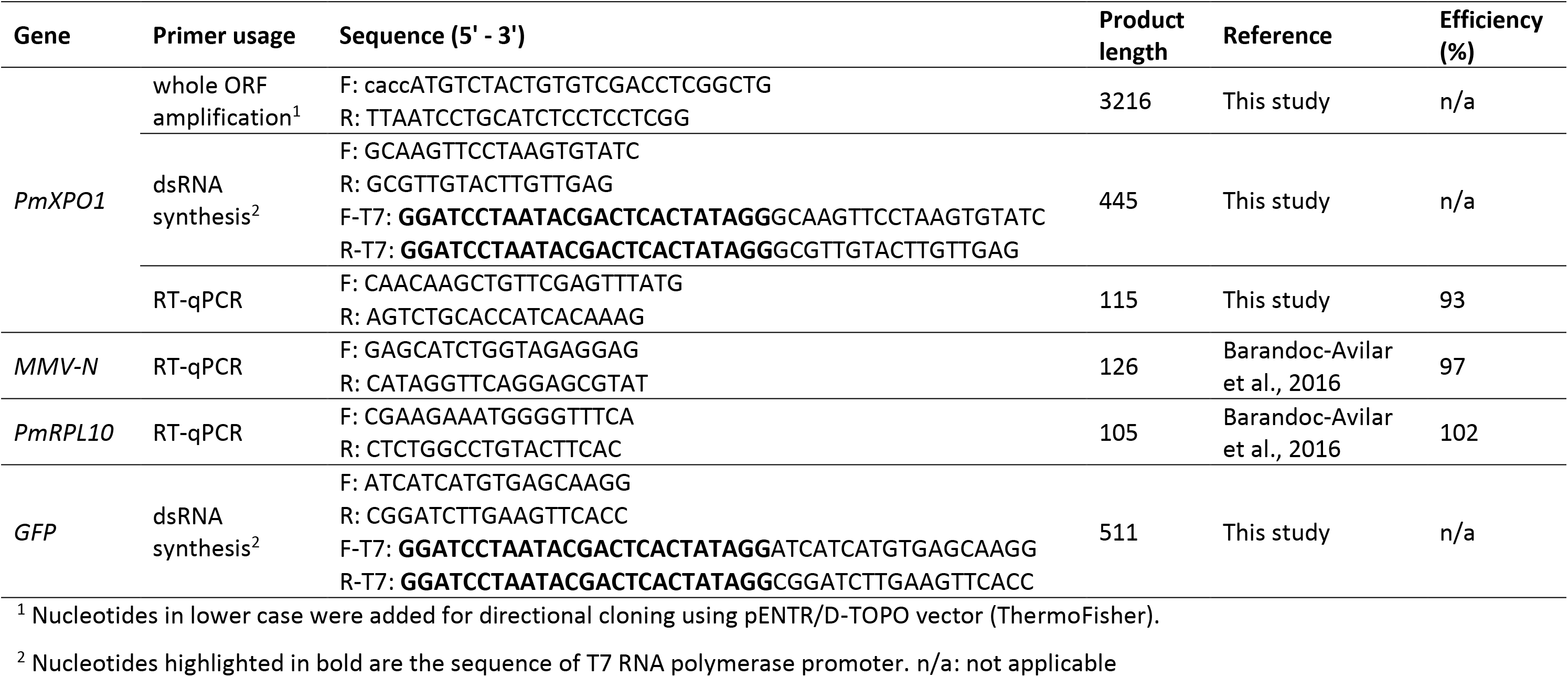
Primers used in this study.

**Supplementary Table S2.** Data set used in phylogenetic and identity analyses.

| GenBank access # | Protein | Organism | Length (aa/nt) | Order |
| --- | --- | --- | --- | --- |
| XP_023704052_1 | exportin_1 | *Aedes aegypti* | 1054/3162 | Diptera |
| KAI_5710562.1 | exportin_1 | *Aphis gossypii* | 1080/3240 | Hemiptera |
| XP_001653392_1 | exportin_1 | *Apis mellifera* | 1063/3189 | Hymenoptera |
| XP_027853473_1 | exportin_1 | *Bemisia tabaci* | 1054/3162 | Hemiptera |
| XP_006571185_1 | exportin_1 | *Cryptotermes secundus* | 1064/3192 | Isoptera |
| XP_018899831_1 | exportin_1 | *Diaphorina citri* | 1079/3237 | Hemiptera |
| NP_001303316_1 | exportin_1 | *Drosophila melanogaster* | 1064/3192 | Diptera |
| XP_011311556_1 | exportin_1 | *Fopius arisanus* | 1064/3192 | Hymenoptera |
| XP_026287654_1 | exportin_1 | *Frankliniella occidentalis* | 1065/3195 | Thysanoptera |
| XP_039285779_1 | exportin_1 | *Nilaparvata lugens* | 1073/3219 | Hemiptera |
|  | exportin_1 | *Peregrinus maidis* | 1072/3216 | Hemiptera |
| XP_049939496_1 | exportin_1 | *Schistocerca serialis cubense* | 1064/3192 | Orthoptera |
| XP_021929121_1 | exportin_1 | *Zootermopsis nevadensis* | 1064/3192 | Isoptera |
| XP_006712157_1 | exportin_1 | *Homo sapiens* | 1072/3216 | - |
| NP_001030303_1 | exportin_1 | *Mus musculus* | 1072/3216 | - |
| NP_197204_1 | exportin_1A | *Arabidopsis thaliana* | - | - |
| XP_015632087_1 | exportin_1A | *Oryza sativa Japonica group* | - | - |
| NP_001349216_1 | exportin_1A | *Zea mays* | - | - |
| AAO34666_1 | exportin_5 | *Arabidopsis thaliana* | - | - |
| NP_608339_2 | exportin_5 | *Drosophila melanogaster* | - | - |
| NP_065801_1 | exportin_5 | *Homo sapiens* | - | - |
| DAA01277_1 | exportin_T | *Arabidopsis thaliana* | - | - |
| NP_009166_2 | exportin_T | *Homo sapiens* | - | - |

**Supplementary Figure S1.**
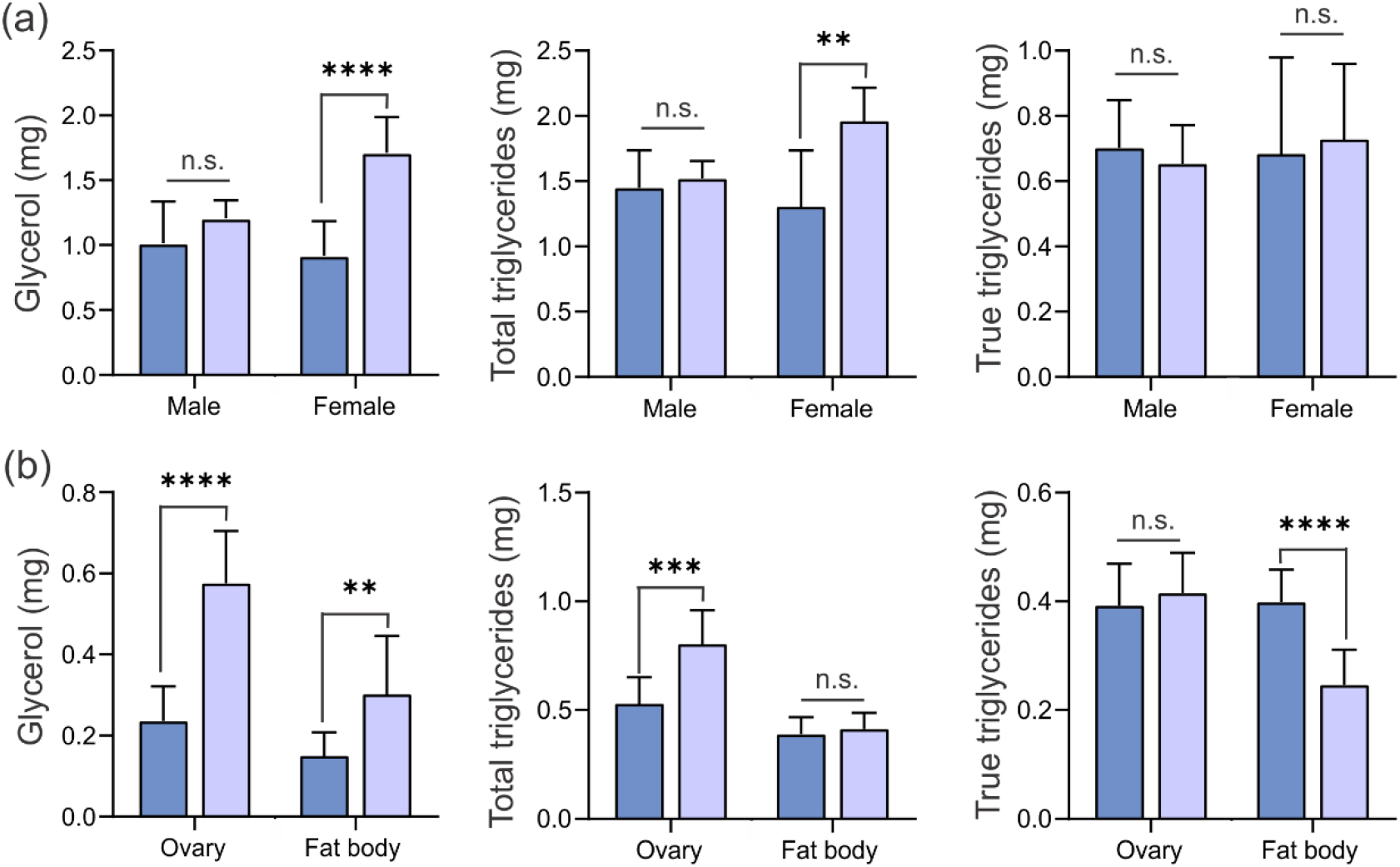
Knockdown of *P. maidis exportin 1* (*PmXPO1*) affects female lipid metabolism. Glycerol, total triglycerides, and true triglycerides were quantified in the whole body of males and females (**a**) and females tissues, ovary and fat body (**b**) at 12 days after dsRNA injection. Bars represent mean and standard deviations of nine and 10 biological replicates for the whole body and tissues, respectively from two independent experiments. Significant differences between groups were assessed by t-test (\*\**p*≤0.01; \*\*\**p*≤0.001; **** *p*≤0.0001; n.s. non-significant).

